# Mitochondrial and Microtubule Defects in Exfoliation Glaucoma

**DOI:** 10.1101/2024.11.25.625249

**Authors:** Arunkumar Venkatesan, Marc Ridilla, Nileyma Castro, J Mario Wolosin, Jessica L. Henty-Ridilla, Barry E Knox, Preethi S Ganapathy, Jamin S Brown, Anthony F DeVincentis, Sandra Sieminski, Audrey M Bernstein

**Author notes:** Address correspondence to: Dr. Audrey Bernstein, SUNY Upstate Medical University Department of Ophthalmology 505 Irving Ave, Syracuse, NY 13210.

## Abstract

Exfoliation Syndrome (XFS) is an age-related systemic condition characterized by large aggregated fibrillar material deposition in the anterior eye tissues. This aggregate formation and deposition on the aqueous humor outflow pathway are significant risk factors for developing Exfoliation Glaucoma (XFG), a secondary open-angle glaucoma. XFG is a complex, multifactorial late-onset disease that shares common features of neurodegenerative diseases, such as altered cellular processes with increased protein aggregation, impaired protein degradation, and oxidative and cellular stress. XFG patients display decreased mitochondrial membrane potential and mitochondrial DNA deletions. Here, using Tenon Capsule Fibroblasts (TFs) from Normal (No Glaucoma) and XFG patients, we found that XFG TFs have impaired mitochondrial bioenergetics and increased reactive oxygen species (ROS) accumulation. These defects are associated with mitochondrial abnormalities as XFG TFs exhibit smaller mitochondria that contain dysmorphic cristae, with an increase in mitochondrial localization to lysosomes and slowed mitophagy flux. Mitochondrial dysfunction in the XFG TFs was associated with an increase in the dynamics of the microtubule cytoskeleton, decreased acetylated tubulin, and increased HDAC6 activity. Treatment of XFG TFs with a mitophagy inducer, Urolithin A, and a mitochondrial biogenesis inducer, NAD^+^ precursor, Nicotinamide Ribose, improved mitochondrial bioenergetics and reduced ROS accumulation. Our results demonstrate abnormal mitochondria in XFG TFs and suggest that mitophagy inducers may represent a potential class of therapeutics for reversing mitochondrial dysfunction in XFG patients.

## Introduction

Age-dependent Exfoliation Syndrome (XFS) is a systemic disease characterized by the formation of aggregated protein deposits called Exfoliation Material (XFM) in all tissues that can synthesize elastin fibers (Nazarali, Damji, & Damji, 2018). Although XFS is a systemic disease, its morbidity is primarily in the eye (U. M. Schlotzer-Schrehardt, Koca, Naumann, & Volkholz, 1992). Exfoliation Glaucoma (XFG) results from protein aggregate formation, leading to a more severe form of Primary Open-Angle Glaucoma (POAG) due to poor prognosis (Ritch, Schlotzer-Schrehardt, & Konstas, 2003). POAG and XFG are associated with high intraocular pressure (IOP), eventually leading to optic nerve damage and blindness. However, XFG displays a marked resistance to classical pharmacological treatments and induces a much faster vision loss (Ritch & Schlotzer-Schrehardt, 2001; Ritch et al., 2003). XFG is strongly dependent on ethnic background, geography, and environmental variables (Pasquale et al., 2014), accounting for 20- 25% of the estimated 70 million worldwide glaucoma sufferers.

The deposition of fibrillar XFM at the Trabecular Meshwork (TM) impedes the outflow of aqueous humor and correlates with increased IOP and visual field loss (Ritch et al., 2003). Several proteomics studies identified multiple proteins in XFM that contribute to the typical XFS/G-related fibril formation, most prominently, extracellular matrix (ECM) cross-linking protein LOXL1, and elastic fiber components such as fibrillin-1, tropoelastin, latent TGF-β binding proteins, microfibril-associated glycoprotein, and clusterin (Morris et al., 2021; Ronci, Sharma, Martin, Craig, & Voelcker, 2013; Sahay, Chakraborty, & Rao, 2022; Sharma et al., 2009). Genome-wide association studies (GWAS) have identified the association of Single Nucleotide Polymorphisms (SNPs) in 8 risk loci with XFG development (Aung et al., 2017; Genetics of Exfoliation Syndrome et al., 2021). Also, an increased propensity of mitochondrial DNA deletions and SNPs in mitochondrial complex I and II genes was reported in a subset of these XFG patients (Abu-Amero, Bosley, & Morales, 2008; Izzotti, Longobardi, Cartiglia, & Sacca, 2011).

XFG is an aging-related disease, and a common finding in aging is abnormalities or decline in cellular fitness, loss of optimal cellular function, and increased cellular senescence (Guo et al., 2022; Miwa, Kashyap, Chini, & von Zglinicki, 2022). Aging worsens protein quality control (David, 2012). Further, many age-related neurological diseases are caused by protein misfolding, the accumulation of toxic aggregates, and the loss of normal proteostasis (Ross & Poirier, 2004). In addition, mitochondrial dysfunction has been increasingly reported in aging and the pathogenesis of various neurodegenerative disorders (Lionaki, Markaki, Palikaras, & Tavernarakis, 2015; Tran & Reddy, 2020). The crosstalk between increased oxidative stress and the accumulation of dysfunctional mitochondria is a hallmark of aging-related neurodegenerative diseases (Lee, Giordano, & Zhang, 2012). Glaucoma shares characteristic features of neurodegenerative diseases and results in progressive loss of optic nerve function (Artero-Castro et al., 2020). The contribution of mitochondrial dysfunction is also strongly implicated in the pathogenesis of POAG patients (Jassim, Inman, & Mitchell, 2021; Vallabh et al., 2022; Vallabh et al., 2024).

Mitochondria are essential organelles for cellular metabolism and survival. They are double-membrane organelles responsible for cellular energy production by oxidative phosphorylation (OXPHOS). However, ATP production also generates a significant amount of mitochondrial Reactive Oxygen Species (ROS) production as a byproduct of ATP biogenesis. In aging-related diseases, mitochondrial function is collectively affected by the accumulation of misfolded protein aggregates, increased oxidative stress and mitochondrial DNA (mtDNA) mutations, compromised organelle trafficking, and impaired elimination of damaged mitochondria by mitophagy (Arduino et al., 2012; Esteves, Gozes, & Cardoso, 2014; Wang, Xu, Musich, & Lin, 2019). Intracellular mitochondrial and other organelle trafficking occurs via the microtubule cytoskeleton. Tubulin acetylation confers mechanical stability for microtubules and dictates organelle transport via motor proteins. However, the deacetylation of microtubules by HDAC6 results in disrupted microtubule arrays and defects in mitochondrial transport (Hubbert et al., 2002).

Despite decades of research on genetic factors, the pathogenesis of XFG is still unsolved. It is becoming increasingly clear that there is a complex molecular interplay behind the pathogenic accumulation of abnormal protein aggregates in the anterior eye segment in XFG (Zadravec et al., 2019). Our previous work demonstrated that XFG TFs have classical aging-related defects, including protein aggregates and dying mitochondria with compromised mitochondrial membrane potential (MMPT) (Bernstein, Ritch, & Wolosin, 2018; Want et al., 2016; Wolosin, Ritch, & Bernstein, 2018). Clinical studies have shown that XFG patients have mitochondrial DNA alterations (Abu-Amero et al., 2008; Izzotti et al., 2011). However, the role of mitochondrial and microtubule dysfunction is mainly unexplored in XFG. Although much experimental evidence indicates that oxidative damage plays a vital role in the pathogenesis of different forms of glaucoma, including XFG, the source of such damage remains to be identified (U. Schlotzer-Schrehardt, 2010). We posit that the loss of mitochondrial function is causative in aberrant proteostasis and an altered oxidative stress response in XFG.

### Materials and Methods Subjects and cell culture

The Institutional Review Board of the State University of New York (SUNY) Upstate Medical University approved this study in compliance with the tenets of the Declaration of Helsinki and HIPAA guidelines. As part of the course of therapy to avoid vision loss, XFG patients often undergo trabeculectomy surgery, in which an incision is made in between the layers of the sclera (white part of the eye) to form an alternative pathway for the aqueous humor to flow. The incision is then “wrapped” with the outer conjunctiva layer and sutured closed. This “bleb” relieves the elevated IOP in the eye. As a part of this surgery, the surgeons remove a piece of tissue called Tenon’s capsule (underneath the conjunctiva) (Promelle, Goëb, & Gueudry, 2021). The tissue can be explanted, and Tenon Fibroblasts (TFs) grow out of the tissue. These patient-derived TFs have served as a productive model system for several labs in the field to understand XFG dysfunction, demonstrating that XFM is released under different stressors (Greene et al., 2020; Want et al., 2016; Zenkel et al., 2011; Zenkel & Schlotzer-Schrehardt, 2014). We receive tissue for no glaucoma (NG) control tenon fibroblasts (TFs) from retinal surgeries and XFG TFs from trabeculectomy surgeries at the SUNY Upstate under an approved IRB protocol. Tenon capsule biopsies are explanted as indicated in our previous publication (Want et al., 2016). Briefly, the tenon capsule tissue samples were explanted over a Matrigel (cat # CB-40230A, Corning) layer in a complete medium, Ham F-12 (DMEM-F12, cat # 11330-032, Invitrogen) complemented with 10% fetal bovine serum (FBS, cat # S11150, R&D Systems) and antibiotic-antimycotic (cat # 15240-062, Gibco). The outgrown fibroblasts from these explants after 5-10 days were used in the following experiments. All subjects were Caucasian, and the deidentified phenotypic data for each donor is listed in Supplemental Table S1.

### Seahorse XFe96 Cell Mito Stress Test

The real-time oxygen consumption rate (OCR) measurement was obtained using a Seahorse Cell Mito Stress Test (Seahorse, Agilent Technologies) on an XFe96 instrument, according to the manufacturer’s protocol. Twenty-four hours before the assay, TF (1 x 10^4^) cells were seeded on an XFe96 cell culture plate (cat # 103793-100, Agilent Technologies) and cultured in DMEM- F12 complete media overnight. On the day of the assay, the cells were washed and replenished with 180 µL of XF assay media (cat # 103575-100, Agilent Technologies) supplemented with 1 mM pyruvate, 2 mM glutamine, and 10 mM glucose. In some experiments, cells were treated with 10 µM Urolithin A (UA) (cat # 6762, R&D systems) or 2 µM NAD^+^ precursor (NR) (cat # 6018, R&D systems) or together for 18 h at the time of cell seeding. During the experiment, the cells were metabolically challenged using 1.25 µM oligomycin (Olig.), 1 µM carbonyl cyanide 4-(trifluoromethoxy) phenylhydrazone (FCCP), and 0.5 µM rotenone plus antimycin A (R/A) (cat # 103015-100, Seahorse XF Cell Mito Stress test kit, Agilent Technologies) for assessing mitochondrial respiration parameters namely, ATP turnover, proton leak respiration, and spare respiratory capacity (SRC). After assay performance, cells were stained with Hoechst dye (cat # 62249, Thermo Fisher Scientific) and measured using Synergy HT BioTek plate reader (BioTek instruments), and the live cell values were used for data normalization.

### Measurement of Mitochondrial ROS

The change in intracellular mitochondrial reactive oxygen species (ROS) was measured by the MitoSOX Red reagent (cat # M36008, Thermo Fisher Scientific). Briefly, TF cells in 24 well plates were incubated with 500 nM MitoSOX Red in Hanks’ balanced salt solution (HBSS) (cat # 24020117, Thermo Fisher Scientific) with calcium and magnesium for 30 min at 37°C. In some experiments, cells were pre-treated with 10 µM UA or 2 µM NR for 18 h before the MitoSOX Red treatment. The probe was removed, and the cells were washed and imaged using a Zeiss LSM 780 confocal microscope with a 63×/1.4 NA oil-immersion objective and live imaging performed at 37°C. The fluorescence signal intensity of mitochondrial ROS was measured by thresholding and normalizing to the cell area using FIJI (1.53 t) software (NIH).

### Mitophagy Assay

To determine the mitophagy flux, NG and XFG TFs were transduced with the mitophagy reporter *Cox8*-EGFP-mCherry retroviral plasmid (Addgene, cat # 78520). This reporter plasmid consists of a *Cox8* mitochondrial targeting sequence at the N-terminus of the EGFP-mCherry fluorescent fusion proteins. While the healthy mitochondria display a yellow color, the mitochondria in the autophagosome lysosomes display red (mCherry) due to the quenching of EGFP (green fluorescence) (Rojansky, Cha, & Chan, 2016). In some experiments, cells were treated with 100 nM Bafilomycin A1 (BafA1, Cayman chemicals, cat # 11038) for 6-8 h. Cells were fixed with 4% PFA, and mounted samples were imaged using a Zeiss LSM 780 confocal with a 63x/1.4 NA oil-immersion objective microscope. Z stack (0.3 µm step size) images were acquired. The number of degrading mitochondria (red puncta) is quantified with FIJI using mito-QC counter macro (Montava-Garriga, Singh, Ball, & Ganley, 2020).

### Microtubule Dynamics

To determine the microtubule dynamicity, NG and XFG TFs were cultured for 24 h in glass bottom dishes in complete media and transfected with the microtubule plus-end marker End-Binding protein 1 (EB1)-mCherry cDNA (Addgene # 55036) (Akhmanova & Steinmetz, 2008). Live cell videos were recorded on a Zeiss LSM510 scanning confocal microscope equipped with a 63×/1.4 NA objective with spatial and temporal resolutions of 110 nm and 0.5-1.0 Hz, respectively. EB1-mCherry tracked microtubule plus-end measurements, and dynamicity was analyzed by µ-track multiple-particle tracking software in the microtubule plus-ends mode (Jaqaman et al., 2008). To validate the assays some cells were treated with nocodazole (50 μM) for 30 min (Fig. S4).

### Immunostaining

NG and XFG TFs were fixed with 4% PFA for 10 min and permeabilized with 0.1% Triton X- 100 for 5 min at RT. Cells were blocked in 10% normal goat serum in PBS for 1 h and stained with alpha-tubulin (cat # 112241-AP, Proteintech), acetylated tubulin (cat # sc23950, Santa Cruz), and mitochondrial marker, HSP60 (cat # NBP3-05536, Novus Biologicals). Further, the cells were stained with secondary goat anti-rabbit Alexa Fluor (AF) 405 (cat # A-31556, Invitrogen), goat anti-mouse AF488 (cat # A-11001, Invitrogen), and goat anti-chicken AF647 (cat # 103-605-155, Jackson Immuno Research) antibodies. The mounted samples were imaged using a Zeiss LSM 780 confocal with a 63×/1.4 NA oil-immersion objective microscope. Z stack (0.3 µm step size) images were acquired. The acetylated tubulin/α-tubulin intensity was calculated using ImageJ software. For some experiments, cells were seeded on collagen-coated coverslips and serum-starved with supplemented-serum-free media (SSFM) for 24 h before immunostaining with anti-LAMP1 and anti-β-tubulin as previously mentioned (Want et al., 2016).

### HDAC6 activity

HDAC6 enzyme activity was measured using an HDAC6 activity fluorometric assay kit according to the manufacturer’s protocol (cat # ab284549, Abcam). The NG and XFG TF cells were harvested and lysed with 100 µl of HDAC6 lysis buffer and incubated with the HDAC6 substrate in a 96-well plate for 30 min at 37°C in a total volume of 50 µl. The reaction was quenched by adding 10 µl of developer solution for 10 min at 37°C. Subsequently, the fluorescence was measured by a Biotech Synergy 2 (Agilent Technologies) microplate reader equipped with Gen5 3.13 software (A_ex_ = 380 nm, A_em_ = 440 nm).

### Electron Microscopy

NG and XFG TFs were fixed with 2.5% glutaraldehyde and 2% paraformaldehyde, followed by 1% OsO_4_ in phosphate-buffered saline (PBS). Cells were dehydrated through an ethanol series (25% to 100%), embedded with EMBed 812 (Electron Microscopy Sciences), and polymerized at 60°C for 24 h. Ultrathin sections were collected on copper 300 mesh grids and stained with UranyLess EM stain (cat # 22409, Electron Microscopy Sciences) and lead citrate and imaged on a JEM-1400 (JEOL) transmission electron microscope (TEM) with Orius SC1000 CCD camera (Gatan, Inc.). 10 individual cell images per sample were collected. As previously mentioned, the mitochondrial structural parameters were quantified using ImageJ (1.53 t) software (NIH) (Lam et al., 2021). The mitochondrial cristae score is evaluated based on the cristae abundance and form, ranging from 0 (worst) to 4 (best), as previously mentioned (Eisner et al., 2017). Briefly, score 0 defines mitochondria with no sharply defined cristae, 1- greater than 50% of the mitochondrial area without any cristae, 2- greater than 25% of the mitochondrial area without any cristae, 3- many cristae but irregular, 4- many regular cristae.

### Statistical Analysis

Data for individual assays were averaged and presented in the figures as mean ± standard error mean (SEM), with the number of samples in each group given in the figure legends. Statistical comparisons were performed using GraphPad Prism (v. 10) as described in the figure legends. Figures were prepared in GraphPad Prism and Adobe Photoshop. Bootstrapping on the mitochondrial area data (Fig. 2B) was performed in Matlab (R2023b) using the *boostrp* function with 500 samples. 95% confidence intervals (CI) were calculated from the bootstrap histogram. To compare the effects of the disease on all examined functions and structures (Figs. 1-4), each data set was range normalized (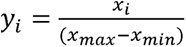, where *x_i_* are the observed values and *x_max_* and *x_min_* are minimum and maximum observed data values from combined patient and control observations for that function or structure), averaged and plotted with coefficient of variation (CV, Fig. 6). Correlation analysis (Supplemental Fig. S5) between the various data sets was performed using *proc corr* in SAS Studio (3.82).

**Figure 1:**
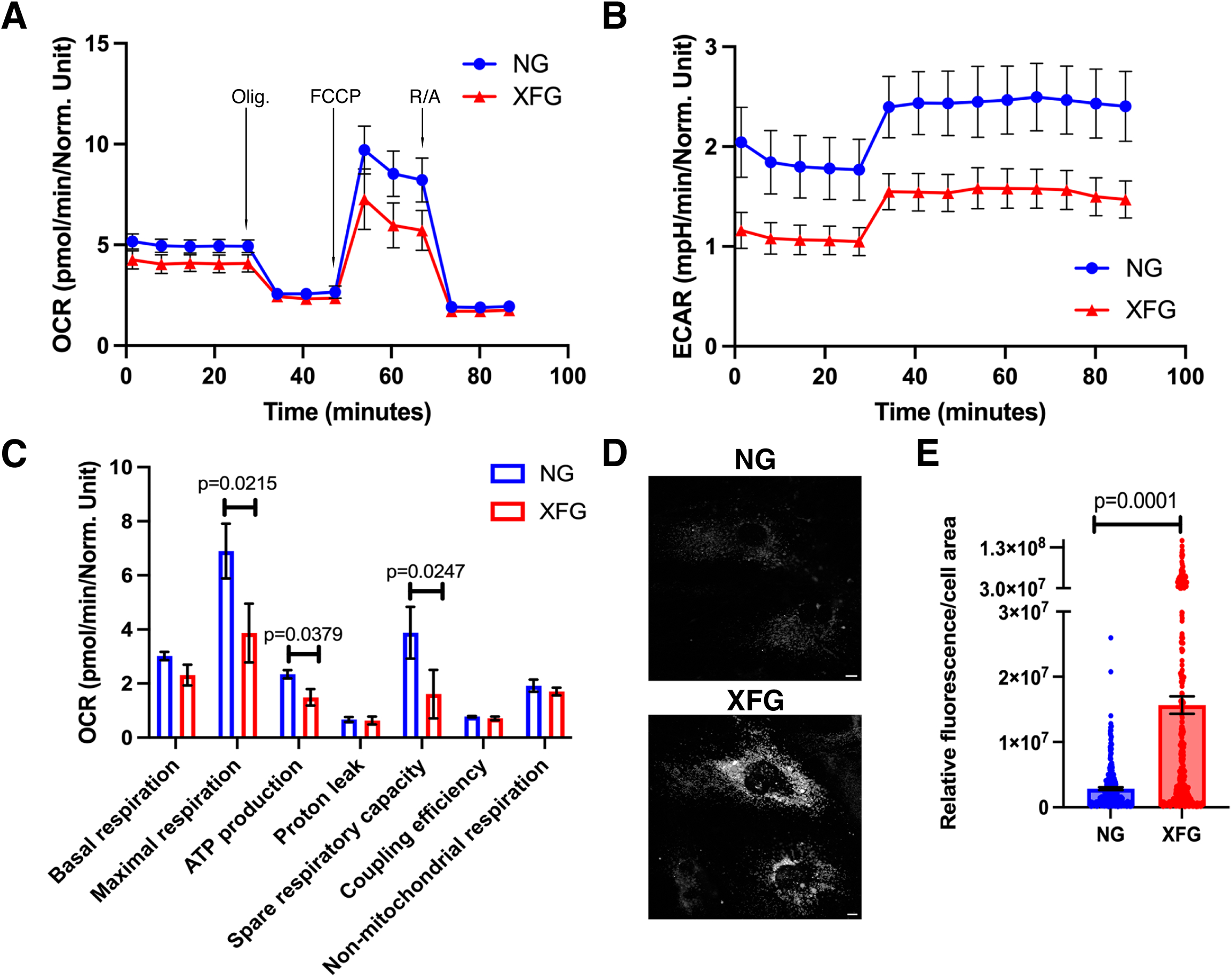
Mitochondrial Bioenergetics is impaired with increased ROS in XFG-TFs. (A-C) Profiles of the Mito Stress Test detected altered mitochondrial cellular bioenergetics for OCR (A) and ECAR (B) in primary NG-TFs (n = 8) and XFG-TFs (n = 7). These data represent a minimum of eight to twelve technical replicates for each primary cell line. Arrows in panel A indicate injections of specific stressors Olig., FCCP, and R/A. (C) Quantification of basal, maximal, ATP-like respiration, proton leak, SRC, coupling efficiency, and non-mitochondrial respiration. Data are normalized by counting Hoechst dye-stained cells (for nuclei) using BioTek Cytation using XF cell imaging and counting software and normalized in Wave software. Data (mean ± SEM) were analyzed using a two-tailed t-test with Welch’s correction. (D-E) Mitochondrial ROS levels were significantly increased in XFG TFs. (D) Representative confocal microscopic images of the same-day analysis of NG and XFG TFs stained with red fluorescent MitoSOX dye (scale bar, 10 µm). (E) The relative fluorescence intensity threshold was set consistently between samples and then normalized to the cell area. Data (mean ± SEM) were analyzed using a two-tailed Mann-Whitney test. NG TFs, 6 independent cell lines, 272 cells (n = 6/272), and XFG TFs n = 5/328

**Figure 2:**
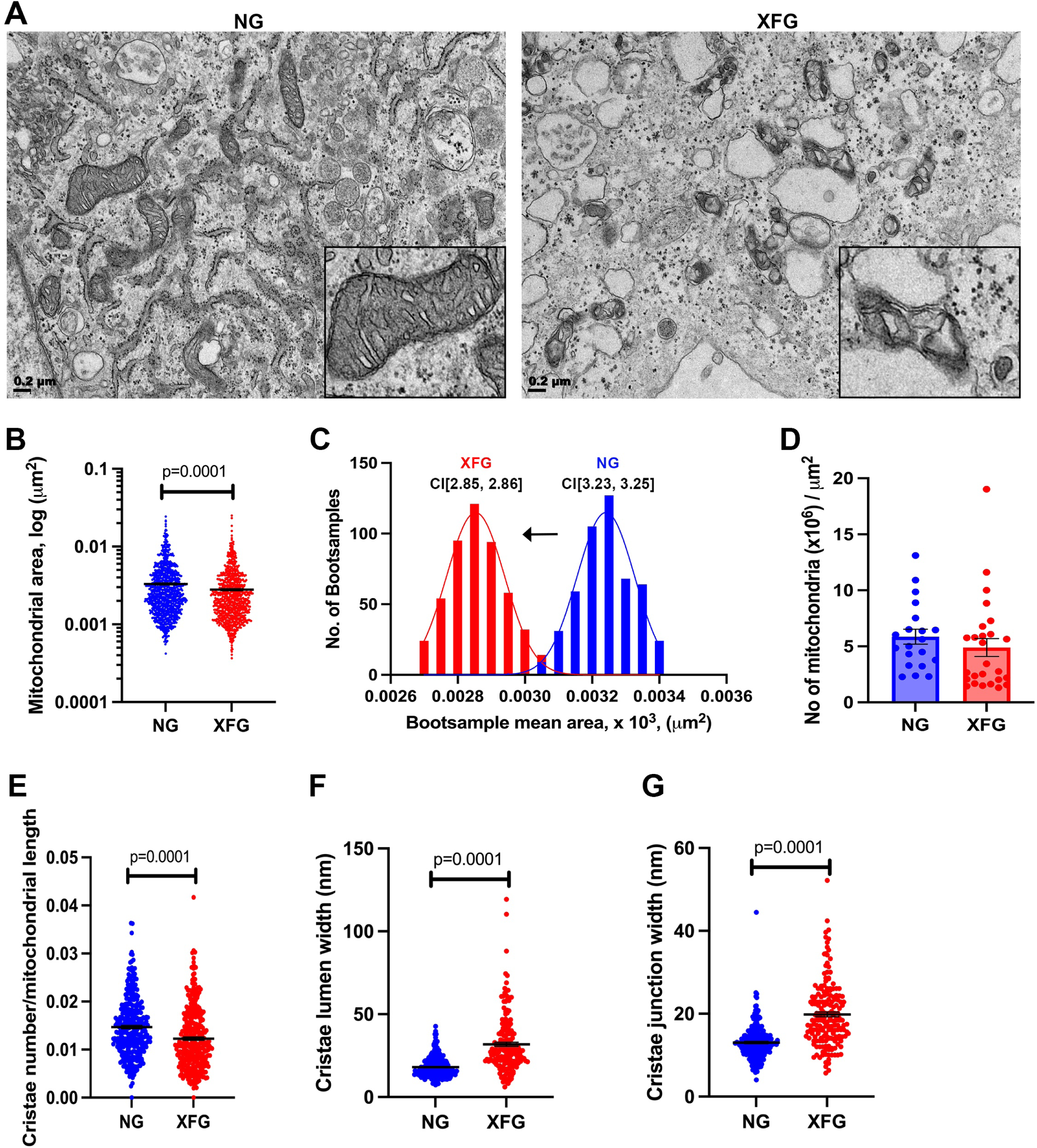
Altered mitochondrial morphology in XFG TFs. TEM of NG and XFG TFs displays smaller mitochondrial organelle morphology with cristae defects from patients with XFG. (A) Representative TEM images of mitochondria from NG (left panel) and XFG (right panel) TFs (scale bar, 200 nm). Boxed regions are shown enlarged inlets. (B) Quantitative analyses of total mitochondrial area per square micron. NG TFs, 3 independent cell lines, 18 cells, 950 mitochondria (n = 3/18/950), and XFG TFs n = 3/25/816. The Y-axis represents the mitochondrial area in the log scale as mean ± SEM analyzed using a two-tailed Mann-Whitney test. (C) The frequency of mitochondrial area from NG and XFG TFs was obtained from TEM images and panel B. Bootstrapping was performed as described in Methods. (D) Quantitative analyses of the number of mitochondria per square micron. NG TFs, 3 independent cell lines, 20 cells (n = 3/20), and XFG TFs n = 3/26. Data (mean ± SEM) were analyzed using a two-tailed unpaired student t-test with Welch’s correction. (E) Quantitative analyses of the number of cristae per mitochondrial length. NG TFs, 3 independent cell lines, 18 cells, 327 mitochondria (n = 3/18/327) and XFG TFs, n = 3/16/342. (F) Quantification of cristae lumen width. NG TFs, 3 independent cell lines, 246 cristae lumen (n = 3/246) and XFG TFs, n = 3/186. (G) Quantification of cristae junction width. NG TFs, 3 independent cell lines, 250 cristae junction (n = 3/250); XFG TFs, n = 3/182. Data (mean ± SEM) were analyzed using a two-tailed Mann-Whitney test (E-G).

## Results

### Impaired mitochondrial bioenergetics in glaucomatous XFG TFs

To assess the possible effects of protein aggregation (U. M. Schlotzer-Schrehardt et al., 1992), defective protein degradation (Hayat, Padhy, Mohanty, & Alone, 2019; Want et al., 2016), and oxidative stress (Mastronikolis, Pagkalou, Plotas, Kagkelaris, & Georgakopoulos, 2022) in XFG on mitochondrial dysfunction, we evaluated the real-time mitochondrial metabolic functions of control NG and glaucomatous XFG TFs by examining their OCR as an indicator of OXPHOS using a Seahorse XFe96 Cell Mito Stress Test. OXPHOS is the primary metabolic pathway through which cells produce the needed ATP from mitochondria. The test showed that the OCR and extracellular acidification rate (ECAR) (an indicator of glycolytic capacity) profiles of the XFG TFs were lower than those of the control NG TFs (Fig. 1A and B), suggesting the overall decrease in cellular respiration in XFG TFs. TFs derived from XFG patients exhibited a significant decline in ATP levels compared to the NG TFs (Fig. 1C; *p* = 0.0379). The decreased ATP production is consistent with the impaired MMPT (Want et al., 2016) as the generated ATP is leaked due to the compromised MMPT. Further, the XFG TF cells also showed significantly lower maximal respiration and SRC than NG TFs (Fig. 1C; *p* = 0.0215 for maximal respiration, *p* = 0.0247 for SRC). These data suggest an impairment of the reserve respiratory capacity in the XFG TFs that would enhance mitochondrial dysfunction in the face of increasing metabolic demand. Thus, XFG disease reveals defective energy metabolism in XFG TF cells.

### Increased oxidative stress in glaucomatous XFG TFs

Mitochondrial OXPHOS is the primary source of mitochondrial ROS generation (Zorov, Juhaszova, & Sollott, 2014). Mitochondrial dysfunction is associated with oxidative stress and is thought to contribute to the pathogenesis of glaucoma (Kamel, Farrell, & O’Brien, 2017). Here, we examined the mitochondrial ROS in XFG TFs using live-cell imaging with MitoSOX Red dye that rapidly accumulates in the mitochondria (Kauffman et al., 2016). We observed an increased fluorescence indicating increased mitochondrial ROS in XFG cells (Fig. 1D). Further, a semi-automated fluorescence intensity analysis showed a significantly higher mitochondrial ROS in XFG TFs than controls (Fig. 1E, *p* = 0.0001). Consistent with the increased mitochondrial ROS in XFG TF cells, we observed an increase in mitochondrial-specific superoxide, SOD2 expression, and reduction in the mitochondrial complex protein, NADH: ubiquinone oxidoreductase core subunit S8 (NDUFS8) using a 2DIGE and mass spectrometry (Fig. S1 and Table S2). Thus, the increased mitochondrial ROS production/accumulation, impaired redox metabolism, and defects in mitochondrial complex protein homeostasis together may be causes for impaired mitochondrial bioenergetics in the XFG TFs.

### Disturbed ultrastructural integrity of mitochondria in XFG TFs

We performed ultrastructural analyses to investigate whether impaired mitochondrial bioenergetics and increased mitochondrial ROS (Fig. 1) in XFG TFs are associated with changes in mitochondrial morphology. Electron micrographs revealed an increased frequency of smaller mitochondria and significantly reduced mitochondrial area (Fig. 2A-C, *p* = 0.0001, Fig. S2A), but without a significant difference in mitochondrial number per cell area in XFG TFs (Fig. 2D).

Bootstrapping approach further confirmed the increase in number of smaller mitochondria in XFG TFs (Fig. 2C, CI[2.85, 2.86] for XFG, CI[3.23, 3.25] for NG).

The mitochondria in XFG TFs showed significantly fewer cristae per mitochondria than in control NG TFs (Fig. 2E, *p* = 0.0001). In addition, while the mitochondria in the NG TFs exhibited subjectively parallel lamellar cristae, mitochondria from XFG patients displayed cristae aberrations such as swollen-, vesicular-, and arc-shaped cristae (Fig. 2A). Also, the mitochondria from the XFG TFs showed a significant increase in cristae lumen area (Fig. 2F, *p* = 0.0001) and junction width (Fig. 2G, *p* = 0.0001). Further, using a scoring system to grade the mitochondrial cristae for their abundance and form ranging from 0 (worst) to 4 (best), the mean cristae score was significantly lower in XFG compared to NG mitochondria (Fig. S2B, *p* = 0.0001). In a comparison of the prevalence of the individual score grades, grades 3 and 4 were more prevalent in the NG TFs, indicating many mitochondria with regular cristae while the score grades were equally distributed in the case of XFG TFs further confirming cristae defects (Fig. S2C). Thus, TEM analysis revealed smaller mitochondria with swollen unstructured cristae for XFG TFs, and these ultrastructural perturbations in the diseased cells may underlie the impaired mitochondrial bioenergetics and homeostasis.

### Impaired mitophagy in XFG TFs

Mitophagy is the process of dead or dying mitochondria fusing with the lysosome for degradation. Because dysfunctional mitochondria are known to degrade selectively through mitophagy, we investigated whether the build-up of poor-quality mitochondria in XFG TFs would lead to an increase in mitophagy using the *Cox8*-EGFP-mCherry mitochondria-targeting pH-sensitive dual emission fluorescent mitophagy reporter containing EGFP and mCherry as fusion proteins (Rojansky et al., 2016). The enhanced green fluorescent protein (EGFP) (p*K*_a_ = 5.9) quenches in acidic compartments, whereas mCherry or red fluorescent protein (RFP) (p*K*_a_ = 4.5) is relatively stable (Ma & Ding, 2021). Cox8 is an inner mitochondrial membrane protein. Therefore, the tandem *Cox8*-EGFP-mCherry targets mitochondria to monitor mitophagy in cells. Normal mitochondria are observed as yellow, whereas autolysosome-enwrapped mitochondria only have a red fluorescence (Ma & Ding, 2021). Following retroviral transduction of the mitophagy reporter in the NG and XFG-TFs, the mitophagy events were observed by red fluorescent puncta on the merged images in Fig. 3A. While the control NG TFs demonstrated fewer red puncta, the XFG TFs demonstrated a greater number of red puncta, indicating that more mitochondria were delivered into acidic lysosomes (mitolysosomes) compared to NG TFs (Fig. 3A). Quantitative analyses showed that the percentage of total mitolysosome area in XFG TFs was significantly higher as compared to the controls (Fig. 3B, NG vs. XFG, *p* = 0.0006). We also quantified the mean size of mitolysosomes and observed a significant increase in mitolysosome size in XFG TFs (Fig. 3C, NG vs. XFG, *p* = 0.0004). The mean mitolysosome size measurements are more reliable than the mitolysosome numbers as the multiple mitolysosomes can fuse together. To address the increase in mitolysosome area and size in the XFG TFs due to the reduced mitochondrial degradation rate in the lysosomes, we analyzed the mitophagy flux. The flux reflects the rate at which mitophagy occurs which is determined by calculating the differences in mitolysosome area and size before and after adding the lysosomal degradation inhibitor, BafA1. We observed a significant increase in the mitolysosome area (Fig. 3B, NG vs. NG BafA1, *p* = 0.0091) and size (Fig. 3C, NG vs. NG BafA1, *p* = 0.0115) only in the NG TFs treated with BafA1. In the XFG TFs, there was not a significant difference ± BafA1, demonstrating that compared to NG TFs, there was no change in XFG TFs ± BafA1, the flux was already defective. This was further confirmed by calculating the 95% CI of the group means between the treatments (Fig. 3B and 3C, right panels). Moreover, TEM also showed numerous abnormal mitochondria in XFG TF cells that are often fused with autophagic vacuoles (Fig. 3D), suggesting mitophagy dysfunction in XFG pathogenesis.

**Figure 3:**
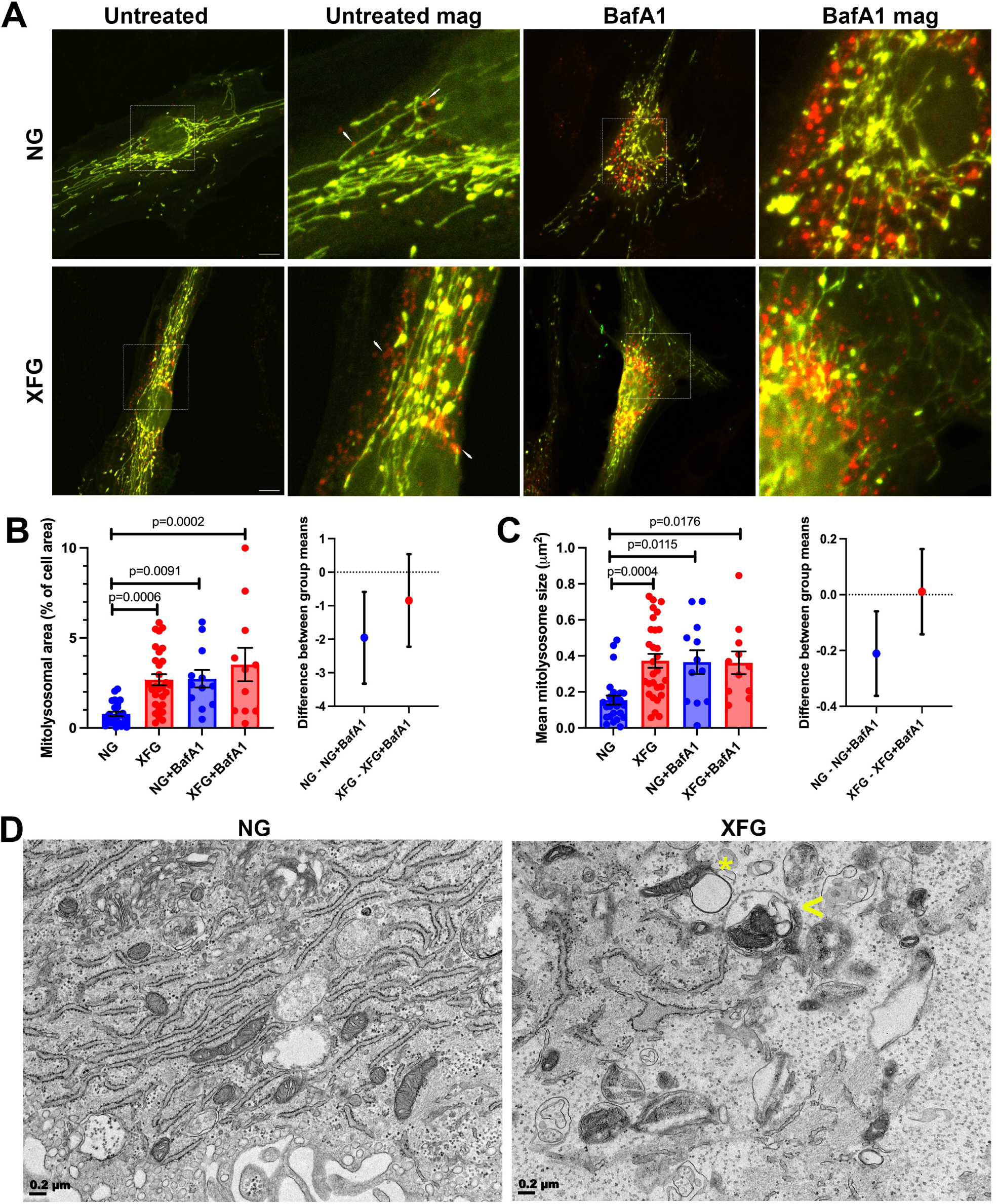
Increase in mitolysosomes in the XFG patient TFs. (A) Representative confocal images of mitophagy reporter (*Cox8*-EGFP-mCherry) signals observed in untreated and BafA1 treated NG and XFG TFs (fixed cells; scale bar, 10 µm). The mitochondria are shown in greenish yellow (EGFP + mCherry), while the degrading mitochondria in lysosomes are shown in red (mCherry) and indicated with white arrows. (B) Quantification of the percent mitolysosome area normalized to the cell area from panel A (left panel). CI plots (95%) of group mean differences of mitolysosome area in the right panel. (C) Quantification of mean mitolysosome size from panel A (left panel). CI plots (95%) of group mean differences of mitolysosome size in the right panel were calculated using Sidak’s multiple comparisons test. Data (mean ± SEM) were analyzed using ordinary one-way ANOVA with Tukey’s multiple comparisons test. NG untreated TFs, 4 independent cell lines, 25 cells (n = 4/25); NG BafA1 TFs, n = 2/12; XFG untreated TFs, n = 4/29; XFG BafA1 TFs, n = 2/11. (D) Representative TEM images of TFs from NG and XFG donors (scale bar, 200 nm). Mitochondria fused with autophagic vacuoles are indicated with *. An arrowhead indicates mitochondria inside the autophagosome-lysosome vacuoles.

### Increased HDAC6 activity and reduced **α**-tubulin acetylation contribute to microtubule hyper-dynamicity in XFG TFs

Mitochondrial dysfunction and mitophagy failure can occur from improper organelle transport along the microtubule network, which provides primary tracks for the trafficking of autophagosomes and mitochondria in the cytoplasm (Melkov & Abdu, 2018). In our earlier work, we demonstrated that when accelerated autophagy was induced by starvation, lysosomes and autophagosomes in XFG TFs did not undergo canonical accumulation at the microtubule organization center (MTOC), which indeed occurred in the control TFs (Want et al., 2016). The lysosomes were clustered in the cytoplasm of XFG TFs after inducing cellular degradation by serum starvation (Fig. S3) (Want et al., 2016), indicating the traffic impairment.

To correlate our previous finding of defective autophagic clearance with microtubule dysfunction, we focused on studying the microtubule plus-end dynamics using EB1, an end-binding protein (Akhmanova & Steinmetz, 2008). We monitored EB1-mCherry on the plus-ends of microtubules in live NG and XFG TF cells using high-resolution confocal microscopy. We tracked several parameters of microtubule dynamics including dynamicity, elongation rate, plus-end growth, and mean lifetime (Fig. 4A-4D). We found that the XFG TF microtubules had elevated dynamicity (i.e., the ratio of the total time growing at the plus-end of microtubules over the lifetime of the microtubule) and were unstable, with a 1.5-fold increase in dynamicity over NG-TFs (Fig 4A, *p* = 0.0001). A significant increase in microtubule elongation rate (Fig 4B, *p* = 0.0003), plus-end growth (Fig 4C, *p* = 0.0026), and growth time (Fig 4D, *p* = 0.0443) in the XFG-TFs contributes to the increased microtubule dynamicity in XFG TFs. Therefore, we hypothesized that faster growth at the plus ends between catastrophes might promote microtubule growth through an additional stabilizing force, like acetylation. To assess microtubule acetylation, TFs were immunostained for acetylated tubulin and total α-tubulin (Fig 4E). We found that α-tubulin acetylation is significantly reduced in the XFG TFs, although the total α-tubulin levels are unaffected (Fig. 4F, *p* = 0.0001). We next analyzed the HDAC6 activity. HDAC6 is a deacetylase that removes acetyl groups from microtubules, thereby regulating microtubule stability (Hubbert et al., 2002). Strikingly, we found a dramatic increase in the level of HDAC6 enzyme activity in the XFG TFs compared with controls (Fig. 4G, *p* = 0.0087). Thus, the reduced acetylated α-tubulin due to increased HDAC6 may contribute to unstable microtubules, impairing organelle transport and altering mitophagy flux in XFG TFs.

**Figure 4:**
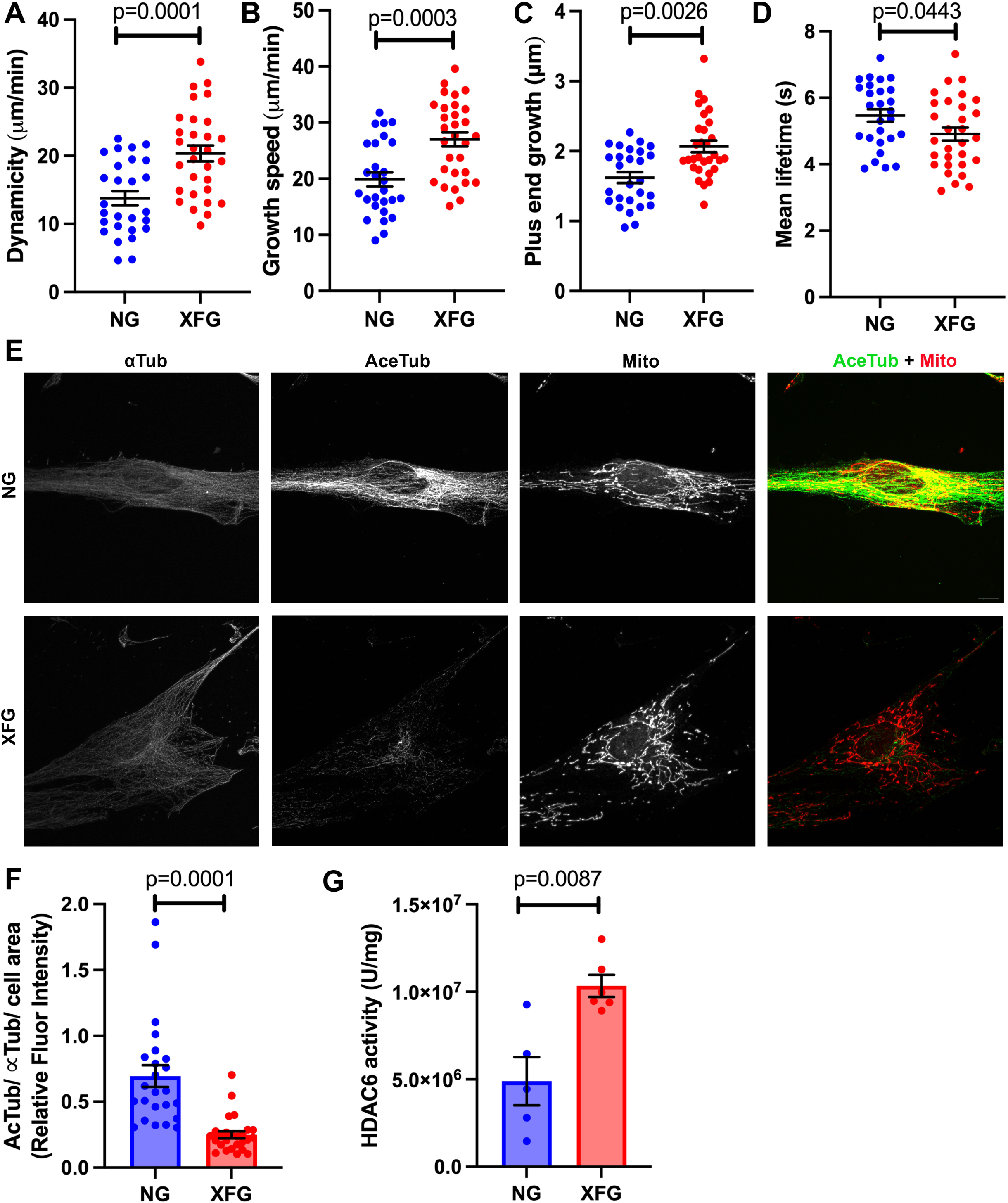
Microtubules are hyperdynamic and less acetylated with increased HDAC6. (A-D) Microtubules are hyperdynamic in the XFG TFs. NG and XFG-TFs were transfected with human EB1-mCherry plasmid, and confocal live cell videos were recorded and analyzed by u-track multiple-particle tracking software in microtubule plus-ends mode (Jaqaman et al., 2008). (A) Quantification of microtubule dynamicity. Dynamicity is an index of overall microtubule dynamics comprising (the sum of all plus-end growth)/(the total lifetime of the microtubule) = time growing plus pauses and catastrophes. (B) Quantification of the average speed of microtubule plus end growth. (C) Quantification of the average length of plus-end growth. (D) Quantification of the average time spent growing. Each point represents one cell and the mean of ∼10,000 microtubule plus-end trajectories from 4 NG and XFG TF independent cell lines each. Data (mean ± SEM) were analyzed using a two-tailed Mann-Whitney test. (E and F) Acetylated tubulin is lower in the XFG TFs. (E) Representative immunofluorescence images stained for α tubulin (αTub), acetylated tubulin (AcTub), and HSP60, a mitochondrial (Mito) marker (scale bar, 10 µm). (F) Quantifying the relative fluorescence intensity of AcTub/ αTub, normalized to the cell area. Data (mean ± SEM) were analyzed using a two-tailed Mann-Whitney test. NG TFs, n = 4 independent cell lines, 24 cells (n = 4/24); XFG TFs, n = 5/26. (G) HDAC6 enzyme activity is higher in XFG TFs. NG (n = 4) and XFG (n = 5) TFs were lysed, and equal amounts of total protein lysates were assayed for HDAC6 activity. Data (mean ± SEM) were analyzed using a two-tailed Mann-Whitney test.

### Urolithin A and NAD^+^ precursor improved mitochondrial health in XFG patient-derived TFs

Using a pharmacological approach to improve mitochondrial health in the XFG TFs, we tested the effects of the mitophagy enhancer, Urolithin A (UA) (Ryu et al., 2016), and the mitochondrial biogenesis inducer NAD^+^ precursor, Nicotinamide Ribose (NR) (Schondorf et al., 2018), in the XFG TFs. First, we measured mitochondrial ROS in cells treated with vehicle, UA, or NR overnight. Treatment of NG and XFG TFs with 10 µM UA or 2 µM NR did not show confounding effects on TF cell viability (data not shown). Both UA and NR significantly reduced ROS generation only in the XFG TFs cells (Fig. 5A and 5B; *p* = 0.0024 for UA and *p* = 0.0001 for NR). Although treating UA and NR slightly reduced ROS generation in the NG TFs, because levels are already low, it is insufficient for statistical significance. Additionally, we measured mitochondrial respiration using a Seahorse XFe96 Cell Mito Stress Test with the XFG TFs. Co-treatment of UA + NR improved the OCR (Fig. 5C) along with the significant increase in basal respiration (*p* = 0.0001), FCCP-induced maximal respiration (*p* = 0.0022), proton leak (*p* = 0.0016), ATP production (*p* = 0.0008), SRC (*p* = 0.0472), coupling efficiency (*p* = 0.0197), and non-mitochondrial respiration (*p* = 0.0010) (Fig. 5D). Also, the co-treatment improved only OCR (Fig. 5C) but not ECAR (data not shown). This suggests that both UA and NR specifically enhance mitochondrial respiration in the XFG TFs but not the glycolysis-linked respiration.

**Figure 5:**
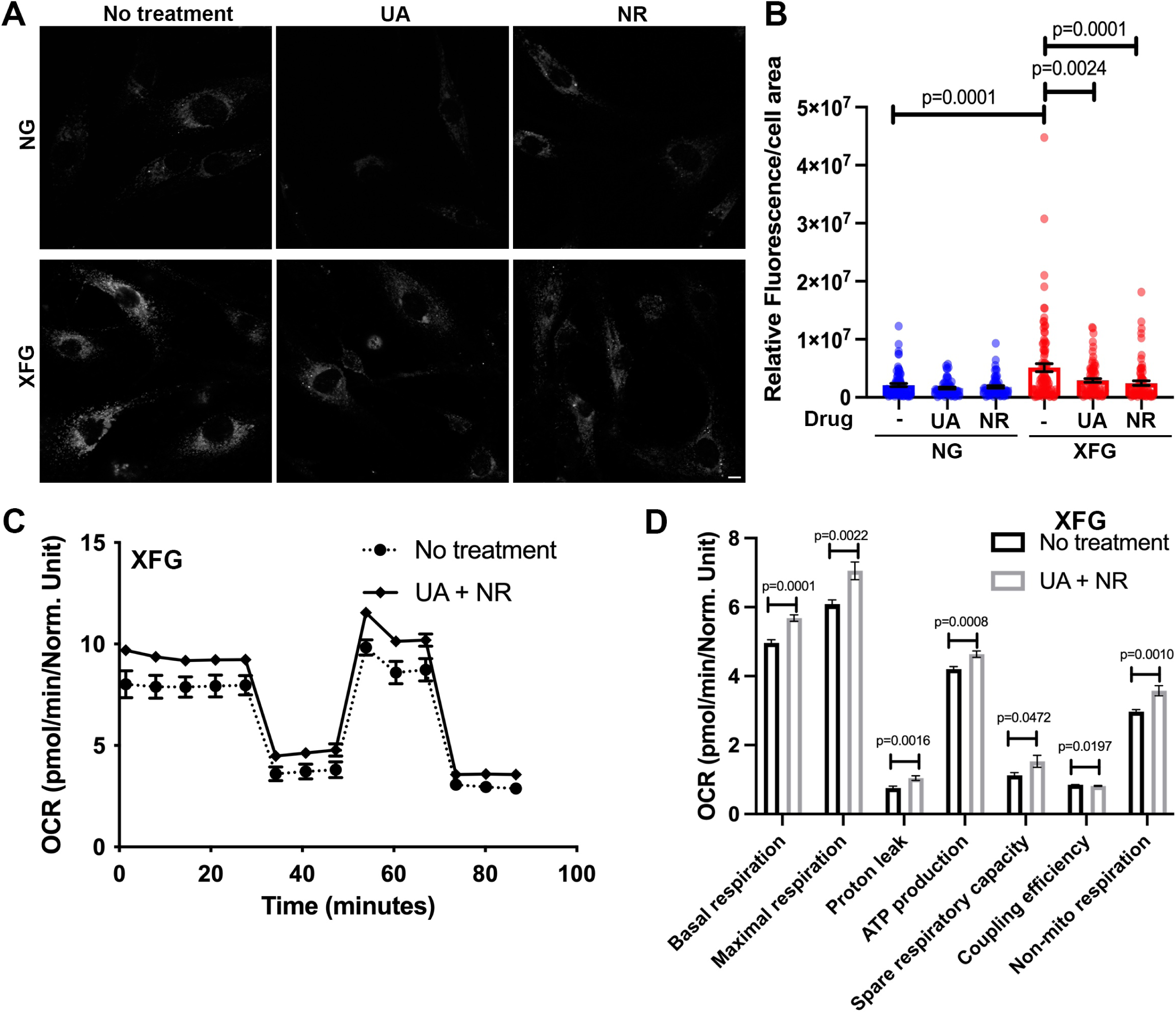
Treatment of UA and NR improves mitochondrial function in XFG TFs. Mitochondrial ROS and bioenergetics were improved in XFG TFs after treatment with mitophagy enhancers UA (10 µM) and NR (2 µM). (A) Representative confocal images of the NG and XFG TFs treated with or without UA or NR overnight and stained with red fluorescent MitoSOX dye (scale bar, 10 µm). (B) Quantifying the relative fluorescence intensity by thresholding and normalizing it to the cell area. Data (mean ± SEM) were analyzed using an ordinary one-way ANOVA with Tukey’s multiple comparisons test. NG TFs no drug, 2 independent cell lines, 80 cells (n = 2/80); UA, n = 2/64; NR n = 2/61 and XFG TFs no drug, n = 2/96; UA, n = 2/76; NR n = 2/75. (C) The maximal OCR was assessed in primary XFG-TFs treated with or without UA and NR overnight (n = 2). These data represent a minimum of eight to ten technical replicates for each primary cell line. (D) Quantification of basal, maximal, ATP-like respiration, proton leak, coupling efficiency, SRC, and non-mitochondrial respiration. Data are normalized by counting Hoechst dye-stained cells (for nuclei) using BioTek Cytation using XF cell imaging and counting software and normalized in Wave software. Data (mean ± SEM) were analyzed using a two-tailed t-test with Welch’s correction.

### Mitochondria, microtubule defects and correlations

Figure 6A compares all the mitochondrial and microtubule properties we measured from diseased XFG and control TFs. To facilitate comparison, since the measurements are very different, the data were scaled using range normalization. It is clear from this plot that there are pervasive deficits in mitochondrial functions, structure, and microtubule dynamics. In fact, correlation analysis documents the relationships between these variables (Fig. S5). This is not unexpected since many of these properties (e.g., basal respiration and ATP levels or cristae junction distance and cristae width) are interconnected. The correlations further support the individual measurements that defined global mitochondrial deficiencies. To examine the size of the effects on each individual property, we examined the percentage change between the XFG and NG TFs. XFG patients exhibited the largest impairments for ROS production (453% increase), mitophagy (267% increase in mitolysosome area and 159% increase in mitolysosome size), and HDAC6 activity (111% increase), with more minor effects in acetylated tubulin, mitochondrial cristae, microtubule dynamics, maximal respiration, and ATP production, basal respiration, cristae numbers, mitochondrial area, and non-mitochondrial respiration (Fig. 6B). These results, in a comprehensive analysis of critical aspects of mitochondria and microtubule functions, demonstrates the deficits observed in XFG patient TFs.

**Figure 6:**
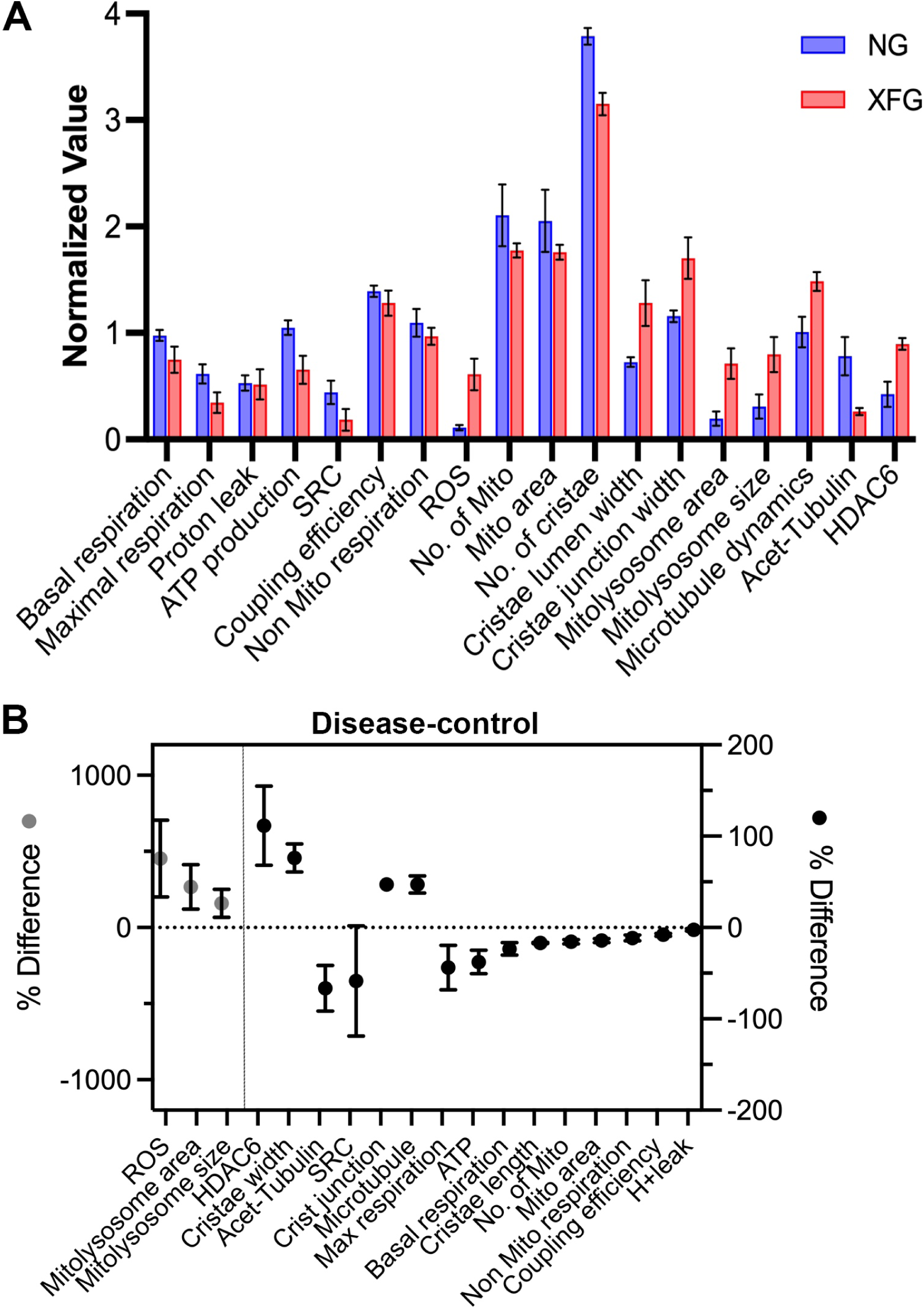
Comparison of the functional and structural effects observed in diseased versus control cells. (A) Data from Figs. 1-4 were range normalized to facilitate comparison of the effect size of XFG. The average difference was plotted with CV (error bars). (B) The average normalized value (XFG-NG samples) calculates the percent difference. The resulting differences are arranged from left to right with decreasing absolute magnitude. The differences in ROS, mitolysosome area, and size are plotted with the left scale, and the remaining differences are plotted with the right scale. Error bars are CV.

## Discussion

A common finding in aging is abnormalities, decline in cellular fitness, and loss of optimal cellular function (Wissler Gerdes et al., 2020). Aging generally declines protein quality control, and many age-related neurodegenerative diseases are caused by protein misfolding and the accumulation of toxic aggregates (Wen et al., 2023). In addition to protein aggregation, there is a well-known age-related dysfunction in mitochondrial dynamics, biogenesis (Lionaki et al., 2015; Lu & Guo, 2020; Srivastava, 2017), dysfunctional microtubule dynamics, aberrant lysosomal positioning, and lysosomal Ca^2+^ signaling (Sferra, Nicita, & Bertini, 2020; Udayar, Chen, Sidransky, & Jagasia, 2022). Together, the build-up of protein aggregates with poor-quality mitochondrial, microtubule, and lysosomal functions underpin many, if not all, age-related diseases. Here, we showed that mitochondrial and microtubule dysfunction in the XFG TFs may have a previously unrecognized role in the pathogenesis of XFG. Using Tenon Capsule Fibroblasts from XFG patients, we have identified defects in mitochondrial bioenergetics, increased mitochondrial ROS production, abnormal mitochondrial morphology, increased mitolysosomes, slowed mitophagy flux, and defective microtubules with increased HDAC6 activity. We also demonstrate that UA and NR positively impact mitochondrial bioenergetics and ameliorate ROS accumulation in the diseased XFG TFs.

The difficulty in studying XFG is the lack of animal models that recapitulate the disease. Because of the identification of LOXL1 as one of the major genetic risk factors, many studies have focused on *Loxl ^-/-^*in the eye. However, these mouse models did not produce XFM or other clinical signs of XFG (Li et al., 2020; Suarez et al., 2023; Wareham et al., 2022; Wiggs et al., 2014; Zadravec et al., 2019). Furthermore, although the *Loxl^-/-^*mice have major systemic abnormalities, including uterine prolapse, aortic aneurysm, enlarged airspaces in the lungs, laxity of skin (Liu et al., 2004), and significant ocular anterior segment structural abnormalities with blood-aqueous barrier disruption (Li et al., 2020; Wiggs et al., 2014), these models failed to produce XFG phenotypes. IOP was not consistently elevated, and these models lacked signs of glaucoma or optic nerve damage (Suarez et al., 2023). One study with mouse *Loxl^+/+^* overexpression in the mouse lens led to transient elevated IOP, but XFM was not generated (Zadravec et al., 2019), suggesting that modulation of *LOXL1* expression is insufficient to recapitulate XFG pathology. Moreover, the influences of mice’s genetic background on *Loxl^-/-^* phenotype further complicate generating an animal model for XFG (Suarez et al., 2023). Interestingly, one model demonstrates characteristics of XFG, the lysosomal trafficking regulator (LYST) knockout mouse with intra-aqueous protein aggregation with pigment dispersion detectable by TEM (Trantow et al., 2009). However, the *LYST* gene is not associated with XFG, although it may suggest a lysosomal contribution to XFG disease. Another study found patterns similar to those seen in the *LYST* rodent after overexpression of *Wnt5a* (Yuan et al., 2019). However, the connection between Wnt5a and XFG is currently unknown.

Thus, to overcome the dearth of appropriate XFG mouse models, we and others have embraced the use of human TFs from XFG patients. Using patient-derived TFs, our group previously showed defects in autophagy, mitochondria, and lysosomal positioning in XFG resembling other neurodegenerative diseases (Bernstein et al., 2018; Want et al., 2016; Wolosin et al., 2018). In addition, other groups have also identified mitochondrial defects in TFs from POAG patients (Vallabh et al., 2022). In support of the TF model system, intraocular POAG primary TM cells and glaucomatous mouse models demonstrate mitochondrial dysfunction due to aging, increased IOP, reduced antioxidant defense, mitochondrial DNA damage, glaucomatous neurodegeneration, and neuroinflammation (He et al., 2008; Izzotti, Sacca, Longobardi, & Cartiglia, 2010; Jassim et al., 2021). Furthermore, aberrant mitochondrial organization was reported in an intermittent IOP stress-induced mouse aging model (Xu et al., 2022). Importantly, however, the clinical significance of mitochondrial dysfunction in XFG disease has yet to be explored.

A striking observation of the present study is that the total cellular bioenergetics are compromised in the XFG patient-derived TFs (Fig. 1). The lack of genes due to the mitochondrial DNA deletions and the SNPs in the complex I and II respiratory chain complex genes could be one of the causative factors for reduced mitochondrial bioenergetics in XFG. Moreover, oxidative stress is commonly observed in glaucoma (Mastronikolis et al., 2022), and the defects observed in mitochondrial bioenergetics (Vallabh et al., 2022) and ROS accumulation in this study are supported by studies in POAG using primary human TM cells from the outflow pathway (He et al., 2008). OXPHOS and ROS production are interlinked processes. Accumulation of ROS in the XFG TFs (Fig. 1D-E) could be due to the altered OXPHOS, altered redox homeostasis, aging, and acquired oxidative stress from environmental factors. Consistent with the increased ROS, the observed upregulation of SOD2 (Fig. S1 and Table S2) (Zenkel, Kruse, Naumann, & Schlotzer-Schrehardt, 2007) in different XFG patient tissue samples suggests the mitochondrial homeostasis is altered and the XFG TF cells redox response is still not enough irrespective of increased SOD2. Also, the altered expression observed with the mitochondrial complex I, NADH: ubiquinone oxidoreductase core subunit family proteins (Fig. S1 and Table S2) (Sahay et al., 2022) along with the SNPs in the complex I and II genes (Abu-Amero et al., 2008) together impact the mitochondrial respiratory functions in XFG. The lower MMPT in XFG TFs (Want et al., 2016) is a collective effect of ROS accumulation and altered complex I genes, causing a leak in the generated ATP from the mitochondrial inner membrane. This suggests that further study elucidating the defects in the whole mitochondrial genome/proteome in XFG pathogenesis is required to advance the understanding of mitochondrial functions.

The excess ROS accumulation is deleterious to mitochondria and other cellular organelles. Similar to our findings in XFG TFs, altered ROS plays a crucial role in the pathogenesis of neurodegenerative diseases (Singh, Kukreti, Saso, & Kukreti, 2019). It is unknown if the increased ROS accumulation in the XFG TFs is the primary or secondary insult to the swollen cristae defects (Fig. 2F and 2G). The decreased mitochondrial respiratory function is consistent with the increased mitochondrial structural defects. The mitochondrial morphological abnormalities observed in the XFG TFs could also occur due to an imbalance between the degradation of defective mitochondria and mitochondrial biogenesis. Mitophagy is a dynamic cellular process that, by selectively degrading the damaged mitochondria, helps to improve mitochondrial function and biogenesis (Lee et al., 2012). The increase in mitolysosomes in the XFG TFs (Fig. 3A-C) implies that defective mitochondria are not efficiently eliminated. However, treatment with the lysosomal degradation inhibitor resulted only in a small insignificant increase in XFG TFs, indicating the reduced mitophagy flux. Hence, the mitochondrial functions are compromised, causing the build-up of cellular stress. It is also possible that the defective mitochondria are targeted for degradation and present in the lysosomal vacuoles but may not be efficiently degraded due to defective transport to MTOC via microtubules. In support of this hypothesis, previous studies in XFG have shown impaired proteasomal (Hayat et al., 2019) and autophagic degradation processes with lower macro autophagic flux (Want et al., 2016).

Microtubules are indispensable for transporting different organelles, including mitochondria and lysosomes, in the cell. Precise organelle positioning is essential for cells to maintain cellular homeostasis, and the integrity of microtubules determines that. Mitochondrial trafficking via the microtubules enables cells to adjust their mitochondrial distribution to changing local needs under different stress conditions. However, microtubule defects can cause impaired transport and localization of autophagosomes and autolysosomes to the MTOC in the perinuclear region, affecting the degradation rate (Mackeh, Perdiz, Lorin, Codogno, & Pous, 2013). In XFG, the lysosomal trafficking was impaired as they failed to localize to the MTOC but not in the NG TFs (Fig. S3) (Want et al., 2016). Using a plus-end growth marker EB1, we observed that the XFG TF microtubule is hyperdynamic (Fig. 4A). The plus ends of the microtubules grow longer and faster and these microtubules exhibited ‘dynamic instability’ transitions. We believe these features ultimately lead to unstable microtubule tracks, a phenotype consistent with other neurodegenerative disorders (Dubey, Ratnakaran, & Koushika, 2015). However, not all the microtubules in the cells at a given time are unstable. Microtubules undergo numerous post-translational modifications, among which tubulin acetylation is one of the factors determining the stability of the non-dynamic microtubules (Akhmanova & Steinmetz, 2008; Mackeh et al., 2013). The reduced acetylated tubulin we observed in XFG TFs (Fig. 4F) corroborates the reduced microtubule stability in the diseased cells. Also, HDAC6 negatively regulates microtubule stability by deacetylating the microtubules (Hubbert et al., 2002). The increased HDAC6 activity observed in XFG TFs (Fig. 4G) correlated with a significant loss of acetylated tubulin and, thus, unstable microtubules. This phenomenon of increased HDAC6 and reduced acetylated tubulin was observed in Huntington’s (Guedes-Dias et al., 2015), Alzheimer’s (Govindarajan et al., 2013), and a neurodevelopmental disorder, Rett syndrome (Gold, Lacina, Cantrill, & Christodoulou, 2015). Treatment with HDAC6-specific inhibitors rescued microtubule stability and mitochondrial trafficking, suggesting the critical role of HDAC6 in neurological diseases (Gold et al., 2015; Guedes-Dias et al., 2015).

In addition to the microtubule defects, the observation that the mitochondria are not well interconnected but are often fragmented in the XFG TFs (Fig. 4E, Mito panel) and their localization mimicking the acetylated microtubules in the cell (Fig. 4E, merge panel) implies that the mitochondrial health is also determined by microtubules and lack of efficient transport may affect organelle health and cellular homeostasis (Cho, Kim, Yu, Kim, & Lee, 2021). In reverse, the defective ATP production and supply also affects the mobility of the motor proteins and causes impaired organelle trafficking. Thus, for efficient cellular homeostasis, both organelles should interactively work together. This has been shown in Parkinson’s disease, where improving the microtubule dynamics and stability has restored axonal transport (Godena et al., 2014) and autophagic clearance (Esteves et al., 2014). Thus, unstable microtubules in XFG TFs due to the hyperactivity of HDAC6 likely negatively affect mitochondrial dynamics, potentially leading to impaired bioenergetics, increased ROS, and accumulation of damaged mitochondria.

Mitochondrial function decreases with aging. Mitochondria-targeted therapeutic strategies are being heavily pursued for metabolic disorders and aging-associated pathologies to improve mitochondrial function (D’Amico et al., 2021). In POAG, mitochondrial-targeted therapeutics are proposed to rescue mitochondrial decline and other pathologies (Kuang, Halimitabrizi, Edziah, Salowe, & O’Brien, 2023). Our present study has shown that UA and NR treatment restores mitochondrial bioenergetics and reduces ROS in the XFG TFs (Fig. 5). Based on our findings, it might be speculated that targeting mitophagy may improve mitochondrial function in XFG patients. UA is a natural compound produced by gut microbiota and has been shown to rescue mitochondrial function in *C. elegans* and mouse models by inducing mitophagy (D’Amico et al., 2021; Ryu et al., 2016). Studies have also shown that replenishing NAD^+^ concentrations is protective against aging-related diseases by enhancing mitochondrial biogenesis and cellular respiration in fly, mouse, and human iPSC models (Saini et al., 2017; Schondorf et al., 2018; Zhang et al., 2016). However, additional work is needed to explain how improving mitochondrial dynamics impacts ECM protein aggregation and clinical severity in XFG patients.

In summary, our present findings indicate that aging, proteostasis dysfunction, mitochondrial and microtubule dysfunction, and oxidative stress affect the cellular physiology of XFG patient-derived cells. Our study represents the first investigation of mitochondrial dysfunction associated with impaired microtubule dynamics in TFs derived from XFG patients. These data reveal that targeting therapeutic pathways based on improving mitochondrial and microtubule function may be effective in slowing the progression of XFG-induced vision loss.

## Acknowledgments

We are grateful to all the donors who contributed to this work. We thank Dr. Xin Jie Chen, a Distinguished Biochemistry and Molecular Biology Professor at SUNY Upstate Medical University, for guidance and relevant discussions. We are grateful to Mohammad Anjum Shaik for his contribution to unbiased TEM analysis. We would also like to thank Dr. Robert Ritch for initiating this cell biology project using patient-derived TFs.

This work was funded by the National Eye Institute (NEI) grants R01 NIH EY024942 and R44 NIH EY035188 to AM Bernstein. Merit Review Award (I01 BX005360) from the United States Department of Veteran’s Affairs to AM Bernstein. The Glaucoma Foundation/Bright Focus Award to AM Bernstein and A Venkatesan. NEI R01 NIH EY030567 to AM Bernstein and JM Wolosin, The Mayer Family Foundation, to AM Bernstein and JM Wolosin., R35 NIH GM133485 to JL Henty-Ridilla.

## Authorship contribution

**A. Venkatesan**: Conceptualization, Methodology, Validation, Investigation, Formal analysis, Writing – original draft preparation, review & editing; **M. Ridilla and J.L. Henty-Ridilla**: Conceptualization, Methodology, Validation, Investigation, Formal analysis related to Fig. 4A-D, Writing – review & editing; **N. Castro**: Investigation; **J. Mario**: Writing – review & editing, Funding acquisition; **B. Knox**: Formal analysis related to Figs. 2C, S2B-C and 6; **P. Ganapathy**, **J. Brown**, **A. DeVincentis III**, and **S. Sieminski**: Surgical sample collection; **A. Bernstein**: Conceptualization, Writing – review & editing, Formal analysis, Visualization, Supervision, Project administration, Funding acquisition.

## Data availability

Data will be uploaded to Dryad and made available upon request.

