## Supplemental for "Mitochondrial and Microtubule Defects in Exfoliation Glaucoma"

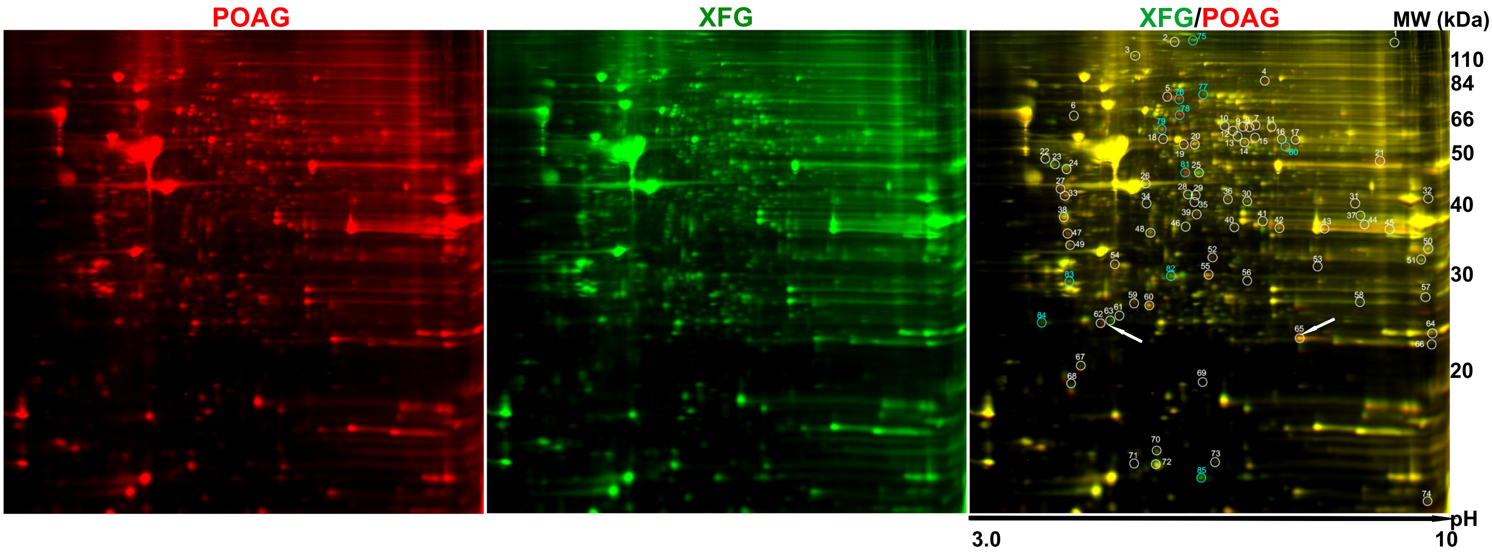


### Figure S1

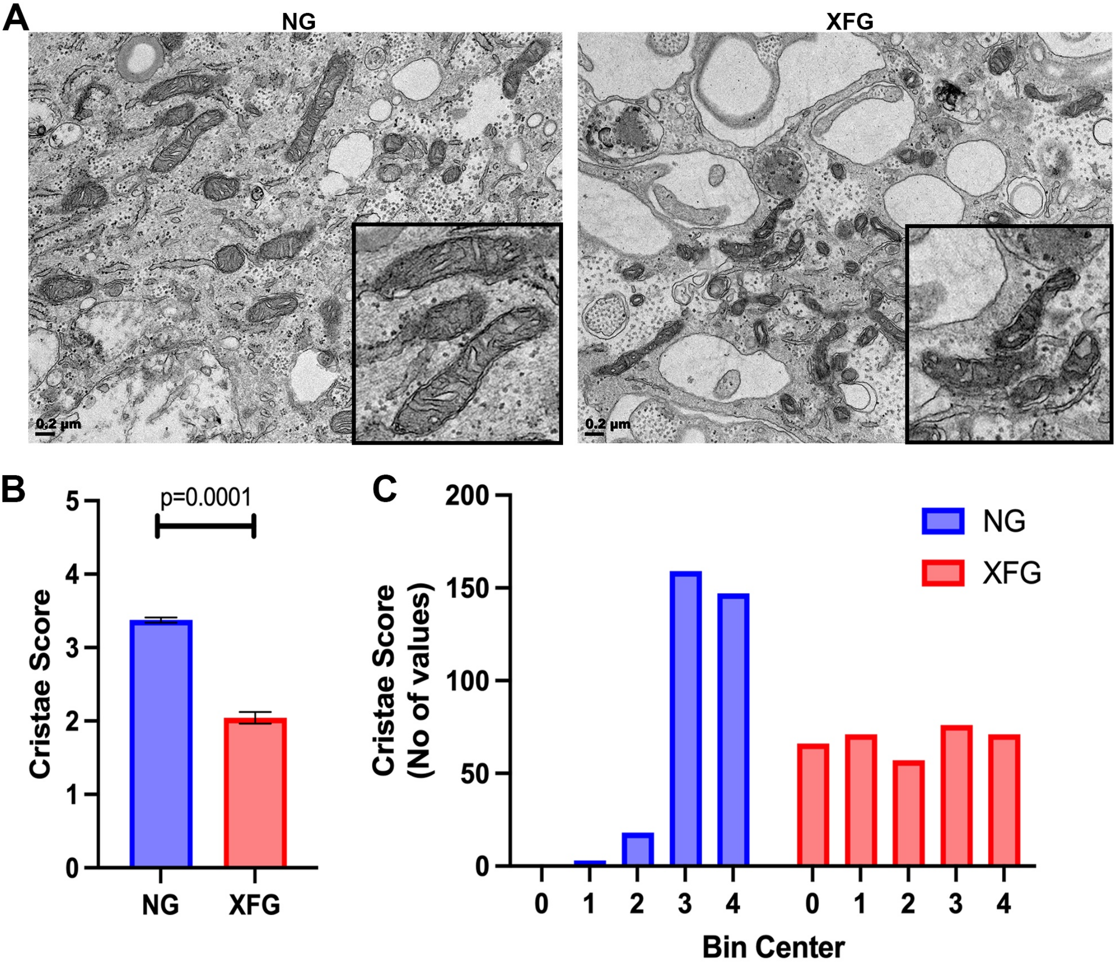

A
NG
XFG
0.2 µm
0.2 µm
B
p=0.0001
Cristae Score
5
4
3
2
1
0
NG
XFG
C
Cristae Score
(No of values)
200
150
100
50
0
NG
XFG
0
1
2
3
4
0
1
2
3
4
Bin Center


**Figure S2**

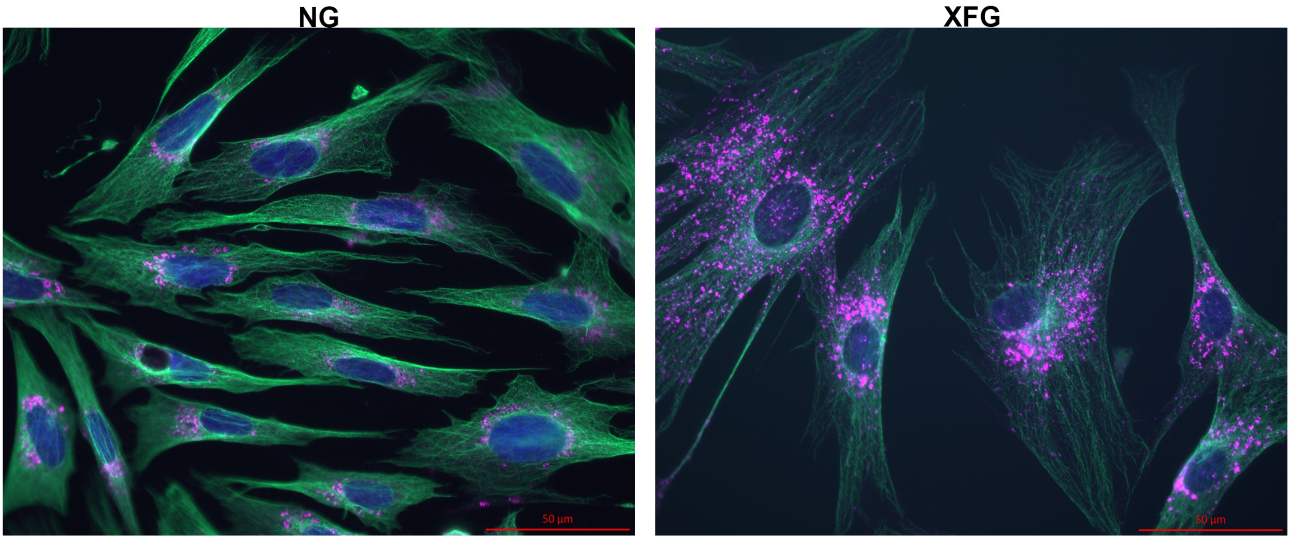


**Figure S3**

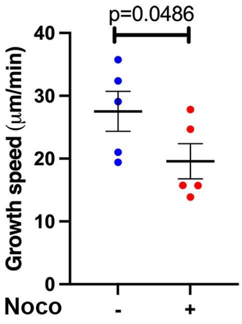


### Figure S4

A

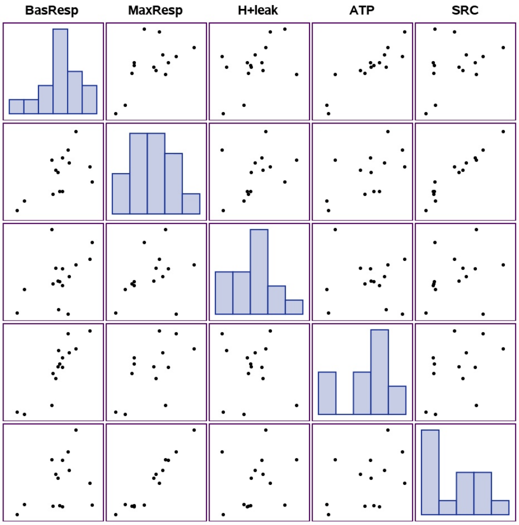


B

| Pearson Correlation Coefficients<br>Prob > \|r\| under H0: Rho=0<br>Number of Observations | | | | | | | |
|---|---|---|---|---|---|---|---|
| | BasResp | MaxResp | H+leak | ATP | SRC | CoupEff | NonMTResp |
| **BasResp** | 1.00000<br><br>15 | 0.59104<br>0.0260<br>14 | 0.47732<br>0.0720<br>15 | 0.89446<br><.0001<br>14 | 0.36432<br>0.2003<br>14 | 0.26461<br>0.3406<br>15 | 0.30702<br>0.2657<br>15 |
| **MaxResp** | 0.59104<br>0.0260<br>14 | 1.00000<br><br>14 | 0.35568<br>0.2120<br>14 | 0.57859<br>0.0383<br>13 | 0.96653<br><.0001<br>14 | 0.17386<br>0.5522<br>14 | 0.35784<br>0.2090<br>14 |
| **H+leak** | 0.47732<br>0.0720<br>15 | 0.35568<br>0.2120<br>14 | 1.00000<br><br>15 | -0.09328<br>0.7511<br>14 | 0.25011<br>0.3885<br>14 | -0.53600<br>0.0394<br>15 | 0.26431<br>0.3411<br>15 |
| **ATP** | 0.89446<br><.0001<br>15 | 0.57859<br>0.0383<br>13 | -0.09328<br>0.7511<br>14 | 1.00000<br><br>14 | 0.42047<br>0.1525<br>13 | 0.62955<br>0.0158<br>14 | 0.19649<br>0.5008<br>14 |
| **SRC** | 0.36432<br>0.2003<br>14 | 0.96653<br><.0001<br>14 | 0.25011<br>0.3885<br>14 | 0.42047<br>0.1525<br>13 | 1.00000<br><br>14 | 0.10740<br>0.7148<br>14 | 0.31565<br>0.2716<br>14 |
| **CoupEff** | 0.26461<br>0.3406<br>15 | 0.17386<br>0.5522<br>14 | -0.53600<br>0.0394<br>15 | 0.62955<br>0.0158<br>14 | 0.10740<br>0.7148<br>14 | 1.00000<br><br>15 | 0.09940<br>0.7245<br>15 |
| **NonMTResp** | 0.30702<br>0.2657<br>15 | 0.35784<br>0.2090<br>14 | 0.26431<br>0.3411<br>15 | 0.19649<br>0.5008<br>14 | 0.31565<br>0.2716<br>14 | 0.09940<br>0.7245<br>15 | 1.00000<br><br>15 |

C

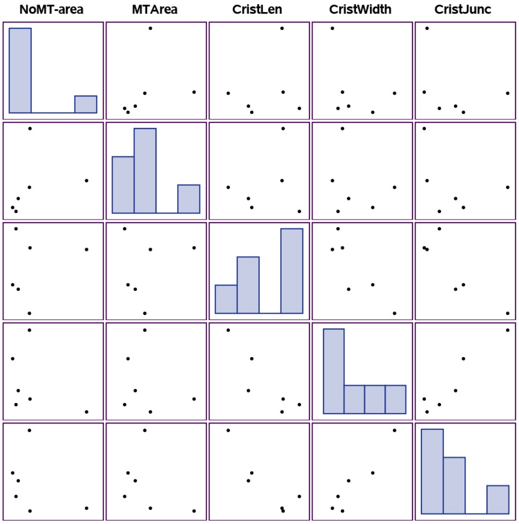


D

| Pearson Correlation Coefficients, N = 6<br>Prob > \|r\| under H0: Rho=0 | | | | | |
|---|---|---|---|---|---|
| | NoMT-area | MTArea | CristLen | CristWidth | CristJunc |
| **NoMT-area** | 1.00000 | 0.28224<br>0.5879 | 0.25134<br>0.6309 | -0.39198<br>0.4421 | -0.36053<br>0.4826 |
| **MTArea** | 0.28224<br>0.5879 | 1.00000 | 0.19797<br>0.7069 | -0.21125<br>0.6878 | -0.37768<br>0.4604 |
| **CristLen** | 0.25134<br>0.6309 | 0.19797<br>0.7069 | 1.00000 | -0.86204<br>0.0272 | -0.87398<br>0.0228 |
| **CristWidth** | -0.39198<br>0.4421 | -0.21125<br>0.6878 | -0.86204<br>0.0272 | 1.00000 | 0.94710<br>0.0041 |
| **CristJunc** | -0.36053<br>0.4826 | -0.37768<br>0.4604 | -0.87398<br>0.0228 | 0.94710<br>0.0041 | 1.00000 |

**Figure S5**

**Table S1:**
Details of the samples used in this study.

| Sample name | Sample source | Age | Sex |
|---|---|---|---|
| NG7 | Retinal surgery | 66 | M |
| NG9 | Retinal surgery | 69 | M |
| NG10 | Retinal surgery | 59 | M |
| NG20 | Retinal surgery | 52 | M |
| NG21 | Retinal surgery | 31 | F |
| NG23 | Retinal surgery | 76 | M |
| NG27 | Retinal surgery | 49 | M |
| NG32 | Retinal surgery | 58 | M |
| NG33 | Retinal surgery | 71 | F |
| NG34 | Retinal surgery | 53 | M |
| NG35 | Retinal surgery | 54 | M |
| NG36 | Retinal surgery | 66 | F |
| NG37 | Retinal surgery | 69 | F |
| NG38 | Retinal surgery | 56 | M |
| NG39 | Retinal surgery | 72 | M |
| NG40 | Retinal surgery | 66 | F |
| NG41 | Retinal surgery | 74 | M |
| NG42 | Retinal surgery | 65 | M |
| NG82 | Strabismus | 4 | M |
| NG83 | Strabismus | 3 | M |
| NG84 | Strabismus | 10 | F |
| Globe NG | Donor Globe | 58 | F |
| X11 | XFG | 67 | F |
| X27 | XFG | 82 | F |
| X44 | XFG | 73 | F |
| XFS002 | XFG | 58 | M |
| XFG21 | XFG | 68 | M |
| XFG23 | XFG | 85 | F |
| XFG24 | XFG | 88 | F |
| XFG26 | XFG | 80 | F |
| XFG27 | XFG | 68 | F |
| XFG28 | XFG | 97 | F |
| XFG30 | XFG | 79 | F |
| XFG32 | XFG | 87 | M |
| XFG33 | XFG | 95 | M |
| XFG34 | XFG | 69 | M |
| XFG35 | XFG | 70 | M |

| | | | |
|---|---|---|---|
| XFG36 | XFG | 78 | F |
| XFG38 | XFG | 89 | F |
| XFG56 | XFG | 89 | M |
| XFG114 | XFG | 77 | F |
| XFG157 | XFG | 70 | M |
| XFG158 | XFG | 74 | M |
| XFG160 | XFG | 82 | F |

NG, no glaucoma; X, XFG, Exfoliation glaucoma. M, male; F, female.

**Table S2: Differentially expressed mitochondrial proteins in XFG-TFs identified by MALDI-TOF/TOF**

| Spot No. | Uniprot No. | Protein name | Gene name | MW(kDa)/ PI | Peptides | Fold change |
|---|---|---|---|---|---|---|
| 65 | P04179 | Superoxide dismutase, mitochondrial | Sod2 | 24.7/8.35 | 9 | 1.8 |
| 63 | O00217 | NADH dehydrogenase iron-sulfur protein 8, mitochondrial | Ndus8 | 23.68/6 | 8 | -2.1 |

*Spot No. is the unique sample number that refers to the labels in Figure S1.
